# Shrimp vaccination with insect-adapted yellow head virus (YHV) extends survival upon YHV challenge

**DOI:** 10.1101/2022.02.13.480220

**Authors:** Warachin Gangnonngiw, Nipaporn Kanthong

## Abstract

This short paper on yellow head virus Type1 (YHV-1) describes preliminary research worthy of further study. YHV-1 disease outbreaks can cause severe mortality in the cultivated shrimp *Penaeus (Penaeus) monodon* and *Penaeus (Litopenaeus) vannamei*. No practical preventative treatment such as vaccination has been reported. However, it has been shown that C6/36 mosquito cell cultures can adapt to YHV-1 and become persistently immunopositive for the virus after 30 split-cell passages or more. Shrimp injection with homogenates from low passages caused yellow head disease (YHD) but from high passages did not, even though injection resulted in immunopositive hemocytes in the injected shrimp. This suggested YHV-1 attenuation during insect cell passaging and the possibility of using cell homogenates as a vaccine to protect shrimp against virulent YHV-1. To test this hypothesis, we injected shrimp with 30th passage homogenates to test for protection against YHD upon subsequent challenge with virulent YHV-1. Results confirmed earlier work that hemocytes of the infected shrimp became both reverse transcriptase PCR positive and immunopositive for YHV-1 but exhibited no mortality. Similarly, there was no mortality in the control group injected with homogenate from YHV-1 negative insect cells. When subsequently challenged with YHV-1, shrimp in the positive control group injected with homogenate from naïve insect cells gave 100 percent mortality within 7 days post challenge while total mortality in the group injected with YHV-1 homogenate did not occur until day 9 post challenge. Kaplan-Meier log-rank survival analysis revealed that survival curves for the two groups were significantly different (p<0.001) and that the mean survival time for the test group (6.5 days) was significantly longer than that in the positive control group (5.4 days). The results confirmed that shrimp injection with YHV-1 immunopositive insect-cell homogenate gave transient resistance to YHV infection and that further research into the possible use of insect cell lines to produce shrimp antiviral vaccines is warranted.

## 1. INTRODUCTION

Yellow head disease (YHD) is caused by yellow head virus (YHV). The virus has been divided into 8 subtypes (Dong et al., 2017; Mohr et al., 2015; Wijegoonawardane et al., 2008a; Wijegoonawardane et al., 2008b), and it has been shown that only Type-1 (YHV-1) and Type-8 (YHV-8) (Dong et al., 2017) cause rapid and severe mortality. This contrasts with 6 less virulent variants (Types 2 to 7), one of which (YHV-2) has been given the specific name of gill associated virus (GAV) (Spann et al., 1997; Spann et al., 2000). *Penaeus (Penaeus) monodon* and *Penaeus (Litopenaeus) vannamei* (the two main shrimp species cultivated in Thailand) are both highly susceptible to YHD caused by two known variants of YHV-1 (YHV-1a and -1b) (Senapin et al., 2010; Sittidilokratna et al., 2009). YHV-8 has not been reported from Thailand.

To date, there are no practical therapeutic treatments available for YHD and the only effective prevention measure has been cultivation of post larvae derived from specific pathogen free (SPF) shrimp in a biosecure setting. Use of dsRNA to knock down YHV non-structural proteins has been shown to inhibit viral replication and lead to improved shrimp survival in laboratory challenge tests (Posiri et al., 2011; Tirasophon et al., 2005; Tirasophon et al., 2007) but no practical applications have yet arisen from such research.

Because there are no immortal cell lines for any crustacean, earlier work was done to determine whether insect cells could be used to maintain and study shrimp viruses (Arunrut et al., 2011; Gangnonngiw et al., 2010). Tests with YHV-1 infectivity in Sf9 lepidopteran cells and in C6/36 mosquito cells revealed that both cell lines became immunopositive for YHV-1 and could be maintained as persistently immunopositive cultures by serial split passage of whole cells. It was found that homogenates of such cells up to 5 split-passages could cause YHD in challenged shrimp, while homogenates from high passages could not (Sriton et al., 2009). Despite the lack of YHD in the shrimp challenged with high-passage YHV-1 homogenates, the shrimp injected with the homogenate did show progressively higher numbers of hemocytes immunopositive for YHV-1 structural proteins post injection (i.e., prior to virulent YHV-1 challenge). This indicated some kind of amplification in the number of immunopositive cells. It was proposed that YHV-1 had been attenuated upon extended split-passaging in the insect cell lines. Similar results were obtained using shrimp white spot syndrome virus (WSSV) (Sriton et al., 2009). The persistent immunopositivity of these cells for many split passages indicated persistent maintenance of the corresponding genes for the relevant YHV-1 and WSSV proteins. Altogether, the results suggested that insect cell lines might provide a mechanism for production of attenuated viruses that could be tested as possible vaccine-like reagents for shrimp.

To test the hypothesis that C6/36 cell homogenates positive for YHV-1 by both PCR and immunohistochemistry might provide protection against YHV-1 in this study, we repeated earlier work to inject shrimp with high-passage YHV-positive or normal C6/36 cell homogenates. We confirmed that the shrimp injected with YHV-1 positive homogenate showed no signs of YHD but did show increasingly YHV-immunopositive hemocytes while those injected with normal C6/36 cell homogenates did not. When the two groups were challenged with infectious YHV-1 to determine whether the YHV-1 positive homogenate provided any protection against YHD, the only beneficial effect was a significant extension in the time before mortality in the groups injected with YHV-1 homogenate. These results were submitted for publication in 2016 but the manuscript was rejected because of its preliminary nature and because the reviewers believed there was no evidence that shrimp had any capacity for specific adaptive immunity. As will be seen in the following paragraphs, evidence for specific adaptive immunity in insects and shrimp now exists. Thus, we have revived our earlier manuscript in the light of this new information should the reinterpreted results be useful in stimulating further investigations towards practical applications for shrimp vaccination against viral pathogens.

Although it was previously believed that shrimp responses to viral pathogens comprise only innate immune responses (Liu et al., 2009), there were early reports of a “quasi-immune” adaptive response to WSSV in *Penaeus japonicus* (Namikoshi et al., 2004; Venegas et al., 2000). In addition, recent work on the mechanism by which insects accommodate viral pathogens has revealed specific, adaptive responses to viral pathogens that are mediated by nucleic acids and a specific RNA interference (RNAi) response rather than antibodies, as occurs in vertebrates. This work has been summarized in a recent review (Flegel, 2020). Similar nucleic acid mechanisms have also been revealed recently in shrimp (Taengchaiyaphum et al., 2021). Although the complete biochemical details for the mechanisms of viral accommodation remain to be revealed in shrimp, results from the work on insects and shrimp suggests that a process similar to vaccination in vertebrates might be possible by administration of “vaccines” comprised of attenuated viruses, similar to the practice commonly applied in vertebrates. However, the mode of protection, instead of occurring via viral protein antigens would be via the nucleic acids that are responsible for production of the shrimp viral proteins detected in cultured insect cells and in shrimp injected with homogenates from those cells.

## 2. MATERIAL AND METHODS

### 2.1 Overall protocol

The overall protocol for the experiments done is shown in **Fig. 1**.

**Figure 1.**
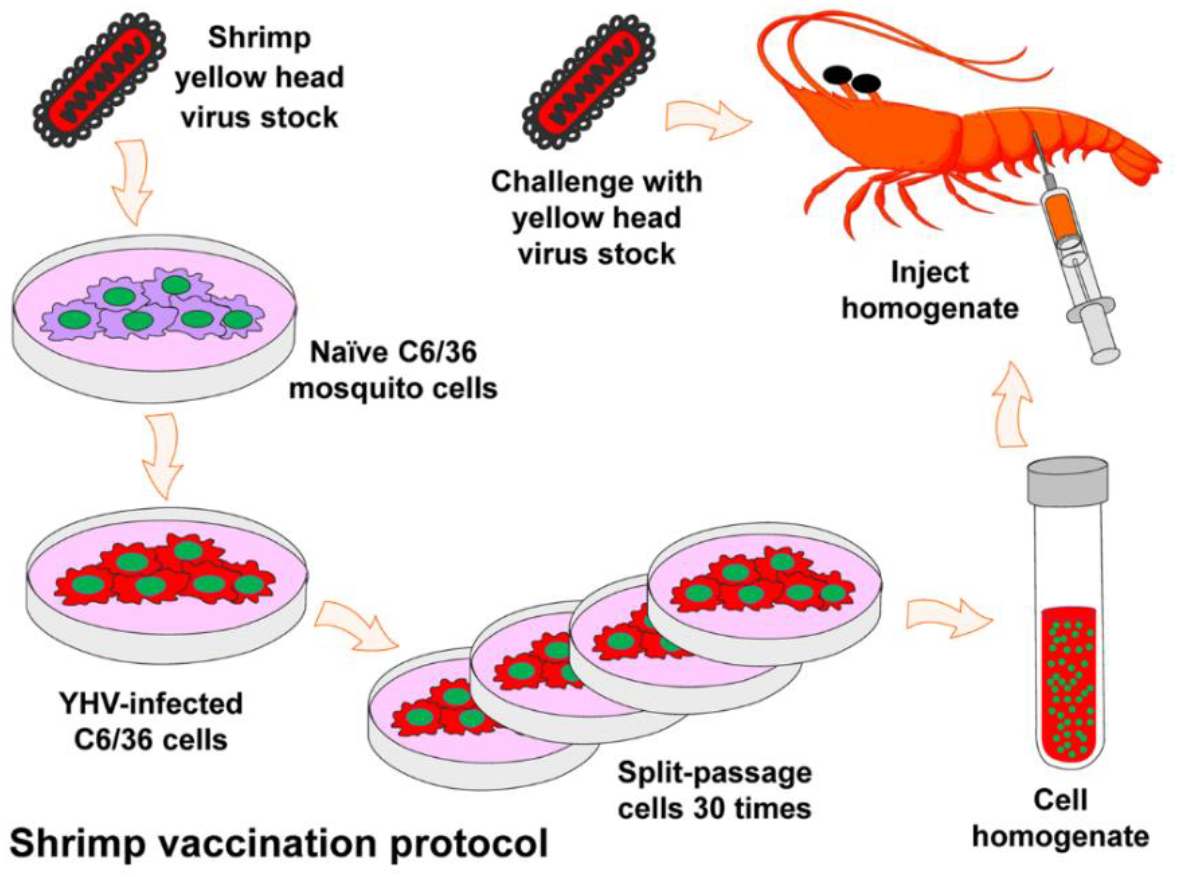
All protocol’s graphic abstract

### 2.2 YHV-1 preparation

Hemolymph of shrimp infected with YHV-1 was collected from the ventral sinus of the first abdominal segment into a syringe containing 10% sodium citrate before dilution with one volume of TNE buffer (0.02M Tris-HCl, 0.4M NaCl, 0.02M EDTA, pH 7.4) followed by centrifugation at 3,000 x g for 30 minutes at 4°C. The supernatant was collected and subjected to ultracentrifugation at 100,000 x g for 1 hour at 4°C. The pellet obtained was re-suspended in TNE buffer and layered onto a 15% sucrose solution (dissolved in TNE buffer containing 20mM Tris-HCl, 1M NaCl, 5mM EDTA, pH 8.0) before ultracentrifugation at 100,000 x g for 90 minutes at 4°C. The pellet obtained was washed with TNE buffer and subjected to ultracentrifugation at 100,000 x g for 1 hour at 4°C and the pellet was resuspended in a small volume of TNE buffer at 4°C. The presence of YHV particles was confirmed by negative staining transmission electron microscopy (TEM). Viral nucleic acid was confirmed by RT-PCR. This stock preparation was stored at -80°C and used to challenge C6/36 mosquito cell cultures and to challenge shrimp in the vaccination tests.

### 2.3 Insect cells

As previously described (Gangnonngiw et al., 2010) C6/36 Mosquito cells (a single cell-type clone obtained from the American Type Culture Collection under catalogue number CRL-1600) were grown at 28°C in Leibovitz’s L-15 medium (Invitrogen, USA) supplemented with 10% fetal bovine serum (Gibco Invitrogen), 10 % Tryptose Phosphate Broth (Sigma) and 1.2 % Antibiotic Penicillin G and Streptomycin (Gibco Invitrogen) in T-25 Flask (Costar, Corning) for 2 days.

### 2.4 Insect cell cultures persistently positive for YHV-1

YHV-1 viral stock (above) was diluted 1:100 with culture medium and used to challenge C6/36 mosquito cells as previously described (Gangnonngiw et al., 2010). Briefly, confluent cells in 25 cm^2^ culture flasks (Costar, Corning) were split 1/5 and grown to confluence in 2 days in a six well culture plate in 1 ml Leibovitz’s (L-15) medium containing 10% heat-inactivated fetal bovine serum (FBS), 10% tryptose phosphate broth (TPB) and 1.2% antibiotic (Penicillin G and Streptomycin). They were then exposed to YHV-1. After incubation for 2 hours with gentle shaking at room temperature, the medium was removed and fresh medium containing 10% FBS was added for further incubation for 5 days at 28°C. For passaging, the supernatant medium was removed and the cells were suspended in 1 ml fresh L-15 medium containing 10% FBS before transfer to a new culture dish at 1:3 split ratio per well. After 2-days incubation (cells reached confluence) the suspension and 1:3 split transfer followed by 2-day incubation was repeated continuously until persistently infected cultures had been established. Mock-infected cells were subjected to parallel split passage to serve as negative controls.

### 2.5 RT-PCR for YHV detection

Cultured insect cells (>10^6^ cells) were centrifuged at 2500 x g. RNA was extracted from pellets with Trizol reagent (Invitrogen, USA) according to the manufacturer’s directions. YHV was determined by RT-PCR using two different protocols, one (the GY2 method) targeting the ORF1b region (amplicon of 135 bp) (Wongteerasupaya et al., 1997) and the other (Y3 method) targeting the overlapping region between ORF1a and ORF1b similar to the target of the IQ 2000 kit (GeneReach, Taiwan) (amplicon 277 bp).

### 2.6 Immunofluorescence detection of YHV in hemocytes of challenged shrimp

After hemocytes were withdrawn from shrimp, they were seeded on cover glasses 15 mm in diameter (Menzel-glaser ^®^, Menzel GmbH & Co KG) in 24 well plates (Costar, Corning). The cells were fixed with 4% paraformaldehyde in PBS for 15 minutes before washing twice with PBS, followed by permeabilization using 0.1% Triton X-100. Cells were blocked by incubation with 10 % fetal bovine serum at 37°C for 1h before exposure to YHV antibody (Y-19) dilution 1:1,000 at 37°C for 1h, followed by washing twice with PBS-T. They were then incubated with GAM Alexa Flour 546 (Molecular probes) dilution 1:500 at 37°C for 1h and washed twice with PBS-T. After counterstaining with TO-Pro 3 (Molecular probes) at dilution 1:500 for 1 h followed by a final wash with PBS-T, a drop of antifade reagent (Prolong^®^ Gold, Molecular Probes) was added and they were covered with cover glasses for viewing with a confocal laser scanning microscope.

Antibodies against the YHV capsid protein p20 (Y19), envelope protein gp64 (Y18) and envelope protein gp116 (V3-2B) (Sithigorngul et al., 2002) were kindly provided by Prof. Paisarn Sithigorngul at Srinakarindwirote University, Bangkok.

### 2.7 Shrimp challenge tests

For time-course incubation activity studies, two preliminary experiments were carried out to determine the appropriate interval to wait after injection of insect-cell homogenates before challenge with YHV-1. Whole cultured insect cells persistently immunopositive for YHV up to the 30^th^ passage and mock infected cells of equal passage number were collected in phosphate-buffered saline (PBS) (approximately 1×10^5^ whole cells/ mL) and homogenized using a sonicator (Vibra Cell) set at amplitude 50 for 30 s. In Experiments 1 and 2, test shrimp (giant tiger shrimp, *P. monodon*, also called black tiger shrimp) were 12-15 g and were obtained from a local shrimp farm in a total batch of approximately 200 shrimp that were acclimatized in the laboratory for 1 day before use in experiments. During acclimatization, a sample of 5 shrimp was taken to test for absence of YHV-1 by RT-PCR. Each experiment employed 3 groups of 25-30 shrimp each. One group comprised the untreated negative control Group A, one the positive control Group B injected with naïve insect cell homogenate followed by YHV-1 challenge (100 µl each of the YHV stock solution diluted 1: 10,000 with PBS buffer) and one the test Group C injected with YHV-1 positive insect cell homogenate followed by YHV-1 challenge (100 µl each of the YHV stock solution diluted 1: 10,000 with PBS buffer). In preliminary Experiment 1, the interval between homogenate injection and YHV-1 challenge was 8 days and in preliminary Experiment 2, it was 10 days. A larger scale Experiment 3 was carried out using the optimum interval between vaccination and YHV-1 challenge (i. e., 10 days) based on the results of Experiments 1 and 2.

For the large-scale test, specific pathogen free (SPF) shrimp (approximately 7 to 10 g each) were obtained from the Shrimp Genetic Improvement Center, Suratthani province and held in 200 L artificial seawater at 15 ppt and at 28°C in a covered, outdoor wet laboratory. They were fed twice daily with a commercial shrimp feed, and excess feed was removed daily. They were acclimatized for 1 d before starting experiments. A total of 325 experimental shrimp were divided into three groups. Shrimp in the negative control Group A (125) were not injected and were not challenged with YHV-1. Shrimp (100) in the positive control Group B were injected (100 µl each) with homogenate from naïve cells while those (100) in test Group C were injected with homogenate from YHV-positive insect cells (100 µl each). After incubation for 10 days all the shrimp remaining in Groups B and C (86 and 78 respectively) were challenged with YHV-1 by cohabitation with 2 shrimp each that had been injected with YHV-1 (100 µl each of the stock virus solution diluted 1:1,000 with PBS buffer). It was estimated that this would cause mortality in the test shrimp within approximately 10 days based on previous experiments. After challenge, shrimp mortality in all 3 groups was monitored, and the experiment ended on day 10 post challenge. Survival curves were statistically compared using Kaplan-Meier log-rank survival analysis with Sigmastat 3.5 software and differences were considered to be significant at p≤0.05.

## 3. RESULTS AND DISCUSSION

### 3.1. Persistent YHV infections were confirmed in C6/36 cells

As previously reported (Gangnonngiw et al., 2010; Sriton et al., 2009), C6/36 cells persistently infected with YHV-1 were successfully produced by serial split passaging of whole cells. Stable cultures 100% immunopositive for YHV-1 were obtained within 2 passages and immunopositive status was maintained for 30 passages. From passage 2 onwards maintenance of YHV-1 was monitored using immunofluorescence for the YHV-1 structural proteins, gp116 and p20. An example photomicrograph is shown in **Fig. 2** for gp 64 only. Similar results were obtained using Mab against YHV p20 and gp116 (not shown). The cells were also positive for YHV by RT-PCR with both detection methods employed (not shown).

**Figure 2.**
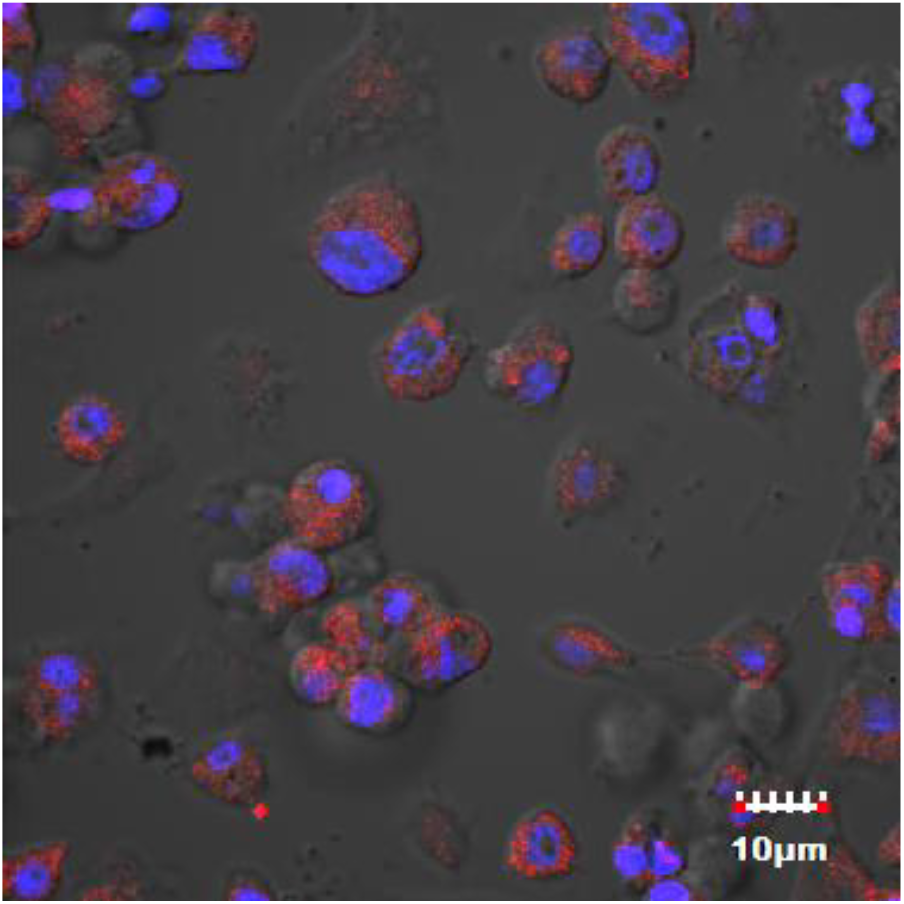
Photomictographs of positive immunofluorescence (red) for YHV in C6/36 cells at passage 22. Red = anti-YHV envelope protein gp 64 protein; Blue = pseudocolor TO-PRO-3 iodide staining of DNA for nuclei.

### 3.2. Confirmation of YHV immunopositive conversion in shrimp hemocytvenes

At 2 days post injection of YHV immunopositive homogenate from C6/36 cells into naïve shrimp, confocal microscopy revealed positive immunofluorescence in hemocytes of the injected shrimp (**Fig. 3**). This immunoconversion was not accompanied by any mortality or gross signs of disease in the injected shrimp. As above for the insect cells, these results confirmed results previously reported for YHV-1 insect homogenates injected into naïve shrimp (Gangnonngiw et al., 2010).

**Figure 3.**
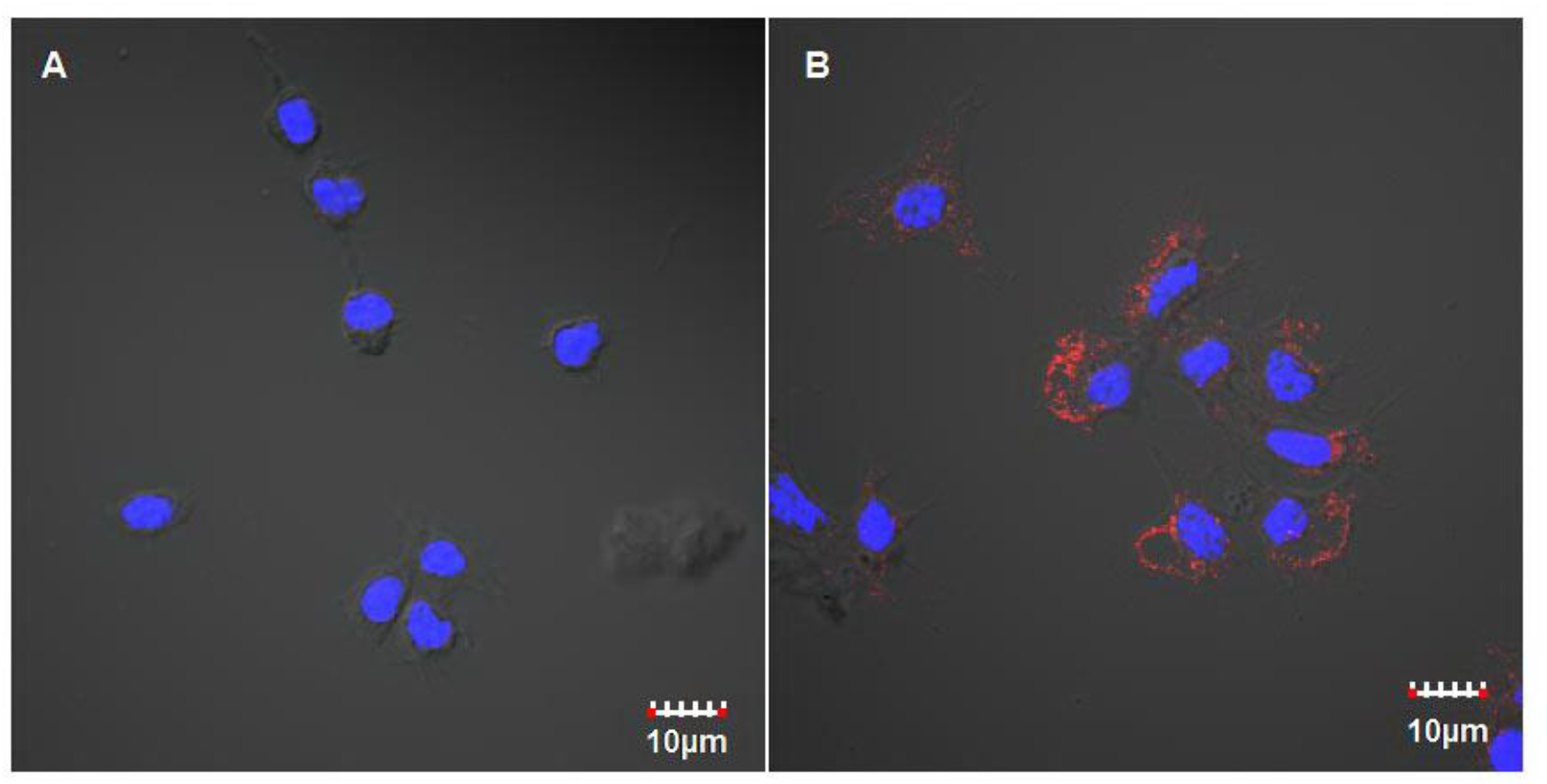
Immunofluorescence photomicrographs of hemocytes from shrimp challenged with homogenates of either (a) whole naïve C6/36 cells or (b) whole yellow head virus (YHV)-immunopositive C6/36 cells from passage 27. The antibody used was anti-p20 capsid protein (red).

### 3.3. YHD mortality was delayed in YHV-immunoconverted shrimp

Previous research on immunization trials against infectious diseases in shrimp required an incubation period ranging from 2 to 15 days after vaccine injection and prior to virus challenge to achieve optimum antiviral effectiveness (Amatul-Samahah et al., 2020). In preliminary Experiment 1 using an interval of 8 days between insect-cell homogenate injection and YHV-1 challenge, the Kaplan-Meier log-rank survival analysis revealed a significant difference (p<0.001) between the negative control Group A and both the YHV-1 challenged groups B (positive control) and C (YHV vaccination). However, there was no significant difference in the survival curve or mean survival time between Groups B and C, indicating no protective effect from injection of the YHV-infected, insect-cell homogenate 8 days prior (**Fig. 4**). By contrast, extending the interval prior to YHV-challenge to 10 days in preliminary Experiment 2 gave a significant difference (p<0.05) between the two YHV-challenge groups with a mean survival time of 8.9 ± 0.6 in YHV-homogenate Group C and 7.4 ± 0.6 in the naïve-homogenate Group B (**Fig. 5**), Thus, an interval of 10 days between injections was chosen for large-scale Experiment 3.

**Figure 4.**
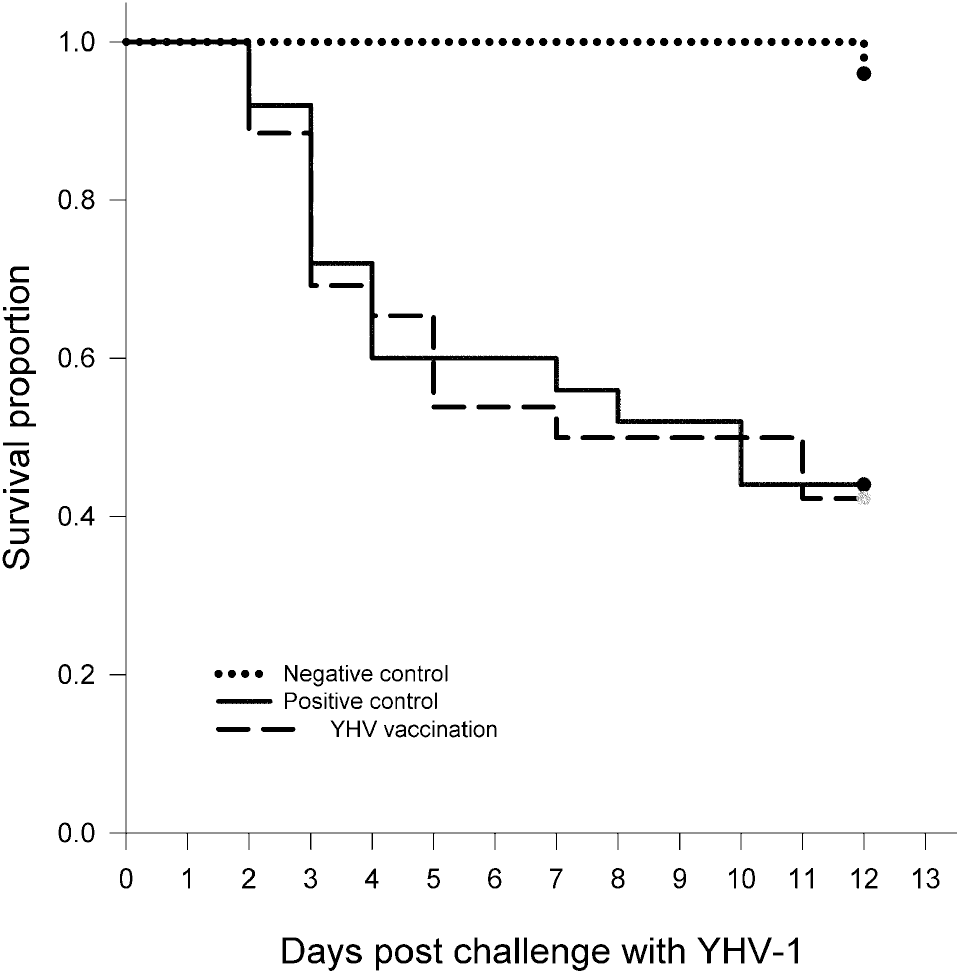
Kaplan-Meier survival curves from preliminary vaccination Experiment 1 using an interval of 8 days between insect-cell homogenate injection and YHV-1 challenge. There was a significant difference (p<0.001) between the survival curve of the negative control group and both of the YHV-challenge groups but no significant difference between the two challenge groups (p>0.05).

**Figure 5.**
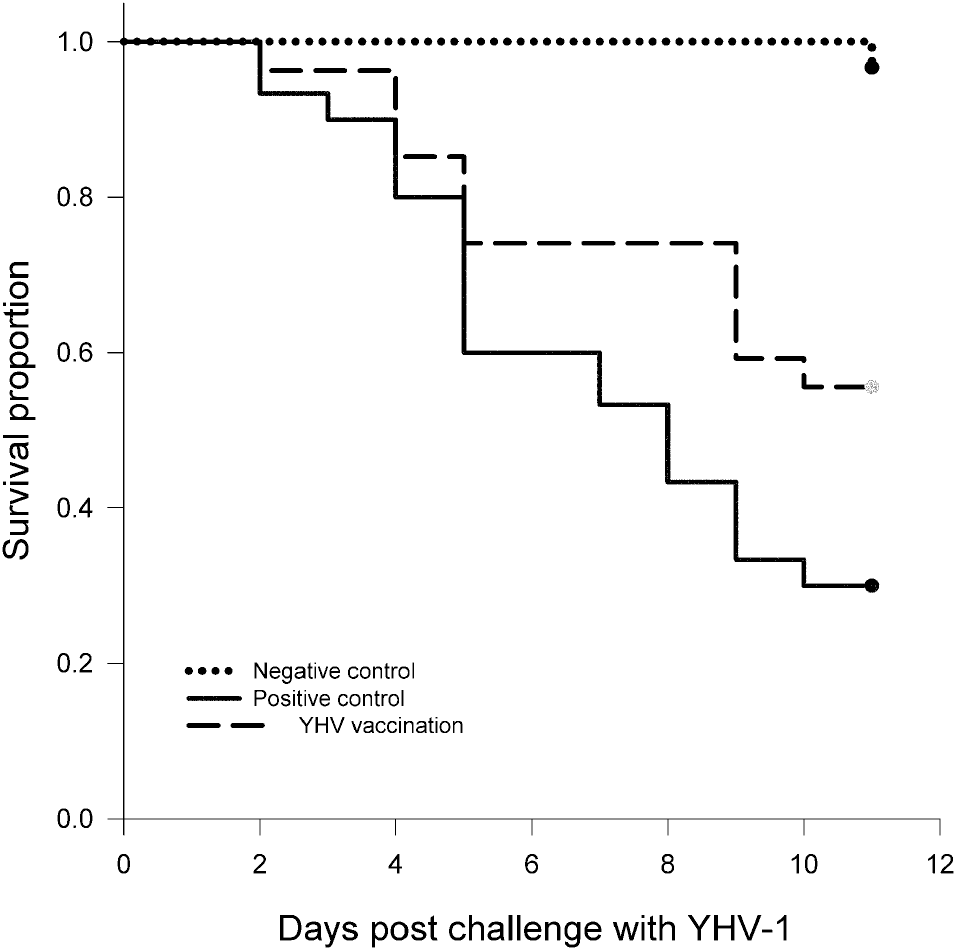
Kaplan-Meier survival curves from preliminary vaccination Experiment 2 using an interval of 10 days between insect-cell homogenate injection and YHV-1 challenge. There was a significant difference (p<0.001) between the survival curve of the negative control group and both of the YHV-challenge groups and also between the two challenge groups (p<0.05).

In large-scale Experiment 3, all the shrimp in the untreated negative control Group A (no YHV-1 challenge) survived. The control test Group C injected with YHV-homogenate showed 77% survival at 5 d post-challenge with YHV-1, compared to only 34% survival in the positive control group injected with homogenate from naïve cells (**Fig. 6**). At day 7 post challenge, survival in Group C was 32% while that in the positive control Group B was zero. By day 9 all the remaining shrimp in test Group C were also dead (i.e., no survival). Kaplan-Meier log rank survival analysis revealed a significant difference (p<0.001) in survival curves and a longer mean survival time (6.5 days) in test shrimp Group C injected with YHV immunopositive insect-cell homogenate when compared to the positive control Group B (5.4 days) injected with naïve insect cell homogenate (Fig. 5). This was similar to the difference in mean survival time (i.e., 2 days) seen in the results from preliminary Experiment 2. Thus, both experiments indicated that injection of homogenates from YHV-immunopositive C6/36 cells was associated with a significant delay in mortality after YHV challenge. Thus, the results from Experiment 3 were similar to those in preliminary Experiment 2.

**Figure 6.**
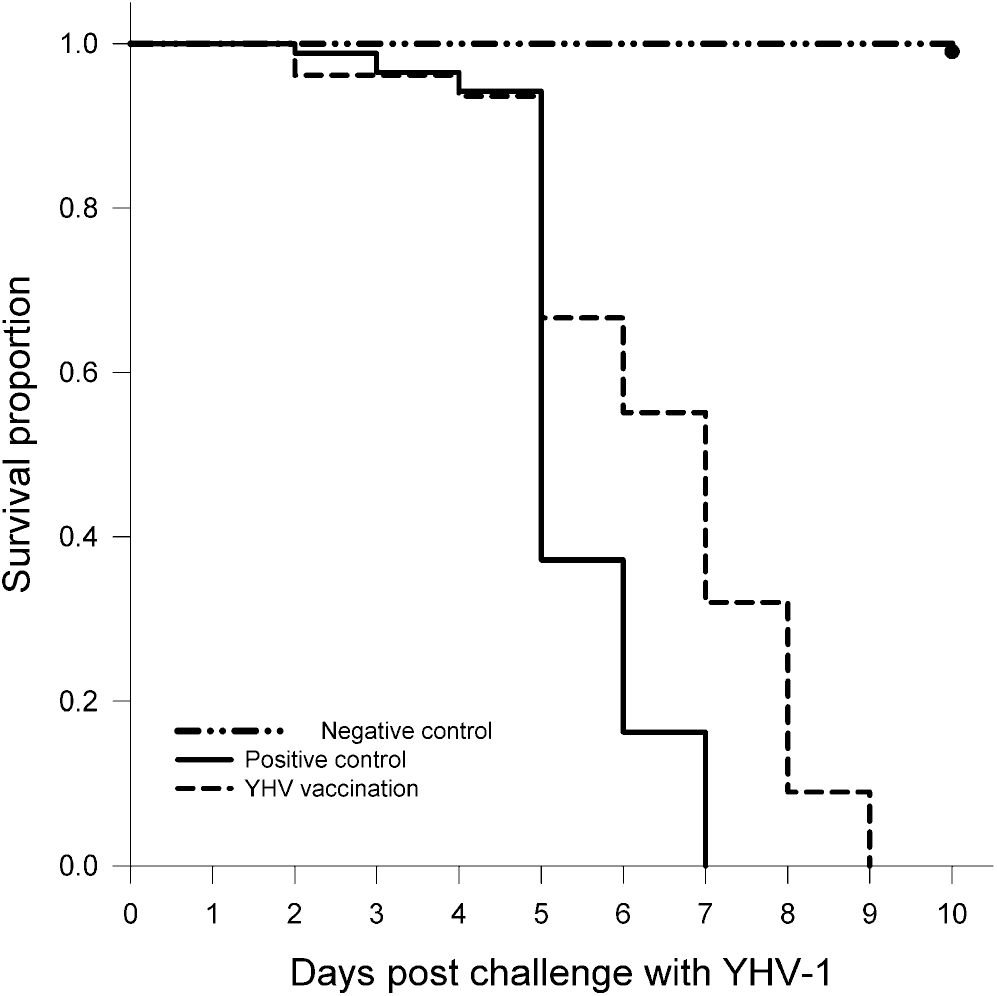
Kaplan-Meier log rank survival analysis results for large-scale Experiment 3, showing a significant difference (p<0.001) in survival curves between the negative control group and both YHV challenge groups and also (p<0.05) between the test groups challenged with YHV-1 where shrimp injected with immunopositive insect cells showed significantly extended survival.

Our hypothesis that prior immune-conversion of naïve shrimp hemocytes to YHV-positive status using homogenates from YHV-immunopositive C6/36 mosquito cells would give protection against mortality upon subsequent challenge with YHV-1 was not upheld. Mortality was 100% in both the test and positive control groups by 9 days in Experiment 3. However, there was a significant difference in the survival curves and mean survival times between the two groups, indicating that the YHV-positive homogenate significantly delayed mortality in test Group C injected with homogenate from YHV-immunopositive insect cells. This was most noticeable on days 5 to 7 post challenge and particularly at day 7 when survival in the positive control group was zero while that in the test group was still 32%. It did not reach zero until almost 2 days later. We believe that these results are encouraging and warrant further tests to determine whether modifications such as lower YHV-1 challenge doses, a modified preparation protocol for the YHV-1 insect-cell homogenate, a longer delay before challenge, or use of booster injections might improve the results. Unfortunately, our project funding expired and our attempts to obtain additional funding to explore alternative experimental protocols were unsuccessful. Our attempts to publish our preliminary results in 2016 resulted in rejection due to their preliminary nature and to lack of evidence that shrimp are capable of a specific immune response.

However, the recent discovery of circular viral copy DNA (cvcDNA) production in response to parvovirus infection in shrimp (Taengchaiyaphum et al. 2021) suggests that the “protective element” in our shrimp study and in the earlier studies on a “quasi-immune” response to WSSV in shrimp may have arisen from viral copy DNA (vcDNA in linear and/or circular forms) in the YHV-1-ommunopositive extracts that were injected into the test shrimp. If this is so, it suggests that insect cell lines such as C6/36 and Sf9 might provide a convenient way to produce vcDNA for shimp viruses *in vitro*. Similar tests previously done with C6/36 cells persistently immunopositive for WSSV (Sriton et al., 2009) should also be repeated to determine whether they produce WSSV-cvcDNA. At the same time, it may be worthwhile testing the same “vaccination” protocol with other viruses such as white spot syndrome virus (WSSV) and Taura syndrome virus (TSV), since they have also been shown give rise to permanently immopositive cells and to yield homogenates that, when injected into naïve shrimp, can immuno-convert their hemocytes but cause no disease (Sriton et al., 2009; Arunrut et al., 2011; Gangnonngiw et al., 2010).

## CONCLUSIONS

Although our experiments did not show that crude homogenates of insect cells persistently immunopositive for YHV-1 would protect shrimp against mortality from challenge with virulent YHV-1, they did show a significant delay in mortality, indicating some degree of induced resistance to YHV-1. Based on current knowledge that specific and acquired antiviral responses of insects and shrimp are mediated by nucleic acids, it is possible that the protective ingredients in the insect cell homogenates was vcDNA, perhaps together with small interfering RNA (siRNA) that may have been produced by the persistently immunopositive cells. We believe that this preliminary work should be followed up to determine whether attenuated shrimp viruses or related nucleic acids produced in insect cell cultures could serve as potential vaccines to protect shrimp, not against viral infections, but against viral diseases.

## Acknowledgements

The authors would like to acknowledge partial support for this work from a research grant provided by Mahidol University and Thailand Research Fund (TRE-CHE grant TRG5680028). They would also like to thank Prof. T.W. Flegel for assistance in editing the manuscript.

